# AKT regulates mitotic progression of mammalian cells by phosphorylating MASTL

**DOI:** 10.1101/375618

**Authors:** Irfana Reshi, Misbah Un Nisa, Umer Farooq, Syed Qaaifah Gillani, Sameer Ahmad Bhat, Zarka Sarwar, Khalid Majid Fazili, Shaida Andrabi

**Affiliations:** Department of Biotechnology, University of Kashmir, 190006 Srinagar, India.; Department of Biochemistry, University of Kashmir, 190006 Srinagar, India.

## Abstract

Microtubule associated serine threonine like kinase (MASTL) has been recently identified as an important regulator of mitosis. By inhibiting protein phosphatase 2A, it plays a crucial role in activating one of the most important mitotic kinases known as cyclin dependent kinase1 (CDK1). MASTL has been seen to be up regulated in various types of cancers and is involved in tumor recurrence. It is activated by CDK1 through its auto regulatory loop but the complete mechanism of its activation is still unclear. In this study, we evaluated the regulation of MASTL via AKT during mitosis. Here we report that AKT phosphorylates MASTL at T299 which plays a critical role in its activation. Our results suggest that AKT increases CDK1 mediated phosphorylation and hence activity of MASTL which in turn promotes cell proliferation.. We also show that the oncogenic potential of AKT is augmented by MASTL activation as the AKT mediated oncogenesis in colorectal cell lines can be attenuated by inhibiting and/or silencing MASTL. In brief, we report that AKT has an important role in the progression of mitosis in mammalian cells and it does so through the phosphorylation and activation of MASTL.

## Introduction

Microtubule associated serine threonine kinase like (MASTL) is known to be one of the main proteins that is essential for mitotic entry in mammalian cells [1]. It is required for timely activation of mitosis by activating CDK1/cyclin B complex via inhibition of protein phosphatase 2A (PP2A) [2]. PP2A is one of the major cellular serine threonine phosphatases that regulates number of signaling pathways [3], as well as cell cylce progression. It is a very complex enzyme and consists of three subunits: the structural A subunit, the catalytic C subunit and the regulatory B subunit, forming a trimeric holoenzyme ABC complex. Each of the three subunits is in turn composed of many isoforms, the most complex being the B subunit. It is believed that the specific combination of the three subunit isoforms imparts specific function to a particular PP2A complex within the cell. The B subunit is the most diverse subunit in PP2A and has very critical role for giving specific localization and function to the enzyme. Among them, PP2A-B55 *δ* isoform is known to be responsible for the dephosphorylation of CDK1 downstream targets to prevent premature mitotic entry [4]. It has been reported that Greatwall kinase, an orthologue of human MASTL, phosphorylates ARPP19 (19 kDa) and endosulphine-*α* (15 kDa, a ligand for sulphonyl urea receptor) at Serine 62 and Serine 67 respectively [5, 6]. After being phosphorylated by Greatwall kinase, these proteins strongly associate with PP2A-B55 *δ*, thus resulting in the inhibition of this phosphatase [6]. Inhibition of PP2A is essential for mitotic entry of mammalian cells, as it facilitates the phosphorylation mediated activation of numerous proteins like CDK1/cyclin B, Plk1, AurK etc. which play critical role in mitotic entry and progression [7]. More evidences have shown that MASTL is necessary for recruitment of cyclin B to anaphase promoting complex (APC) at late metaphase. The timely degradation of cyclin B at this step is necessary for the transition of cells from metaphase to anaphase [8].

In addition to its role at mitotic entry, MASTL plays an important role in mitotic exit as well, where upon its deactivation has been seen to be an essential requirement for the cells to exit from mitosis [9, 10]. Being so critical for mitosis, MASTL has been investigated in various types of carcinomas and has been seen to be up regulated in a number of malignancies [11–13]. Role of MASTL in tumor progression has been attributed to the increased rate of cell proliferation driven by activation/up-regulation of this kinase in tumor cells. Some recent evidences have also shown that MASTL promotes tumor formation and invasiveness via activation of AKT. MASTL has been observed to stabilize S473 phosphorylation of AKT by activating GSK3 which results in the ubiquitination of PHLPP, a phosphatase responsible for dephosphorylation of AKT on 473 [14]. Moreover, some high throughput studies have shown that MASTL can be a better therapeutic target to prevent tumor progression and resistance developed against anti-cancer drugs [11, 15, 16]. Taken together, MASTL regulation is critical for cells both in terms of correct timing of mitotic entry as well as regulation of normal cell proliferation to avoid tumor progression. However, the detailed mechanism of its regulation is still not clear. Although it has been shown that CDK1-cyclin B and more likely Plx1 kinase are responsible for phosphorylating and activating MASTL, but the complete network of its regulation is still not well understood [17].

Here we report that following phosphorylation and activation by CDK1-cyclin B, MASTL is phosphorylated by AKT at T299. This phosphorylation leads to the further stabilization of CDK1-cyclin B mediated phosphorylation and hence activation of MASTL. Moreover, co-expression of MASTL and AKT in HEK293T cells and in SW480, a colorectal cancerous cell line, significantly increases mitotic index of these cells. We also show that the oncogenic potential of MASTL increases substantially in presence of AKT. Together, these results show that MASTL is a bona fide substrate of AKT and the role of AKT in tumorigenesis may largely occur through the activation of MASTL. These results also highlight an unknown role of AKT in the regulation of mitosis in mammalian cells.

## Results

### MASTL is a potential target for AKT

In order to get an understanding of the functions of MASTL, we used Scansite, (www.scansite3.mit.edu), an online computational tool to look for kinases that could potentially target MASTL to regulate its activity. The results indicated that MASTL could be a possible substrate for AKT, as it has a perfect consensus site for AKT phosphorylation (RXRXXS/T) at T299, where R represents Arginine, X any aminoacid, S Serine and T Threonine (Figure S1A). A comparison of the sequences from different mammalian species using ClustalW software (https://www.ebi.ac.uk/Tools/msa/clustalo) showed that this site is highly conserved in many mammalian species (Figure S1B). To directly test whether MASTL is a bona fide substrate for AKT, we co-transfected pCMV-HA-MASTL with increasing amounts of pCDNA3.1-HA-AKT in HEK 293T cells. Results showed that at higher expression of AKT, MASTL protein levels were markedly reduced (Figure 1A-B). This was probably because AKT phosphorylates MASTL at its consensus motif due to which it is subjected to proteasomal degradation. To test whether that is so, we mutated MASTL (Mut-MASTL) at Threonine 299 to Alanine (T299A) (Figure S1C) and co-transfected it with increasing amounts of AKT. Our results showed that mutant MASTL (T299A) protein was not degraded even after the addition of 1.5*μ*g of AKT DNA to HEK 293T cells (Figure 1C-D). To check whether MASTL protein degradation was due to AKT mediated phosphorylation followed by proteasomal degradation, we used bortezomib, an anticancer drug that inhibits proteasomes. Results showed that after the addition of 1*μ*M bortezomib to HEK 293T cells for 24 hours, MASTL protein levels were stabilized back to normal levels (Figure 1E). Furthermore, to confirm that AKT is responsible for MASTL protein degradation, LY294002, a well-known inhibitor of PI3K/AKT pathway was used to inhibit AKT activation in these cells. As can be seen from the Figure 1F, protein levels of MASTL were restored to normal levels by the addition of 20μM LY294002 to HEK293T even in presence of AKT. Moreover, a kinase dead mutant (KD) of AKT (pCDNA3-T7-AKT1-K179M-T308A-S473A) did not affect MASTL protein levels much when transiently expressed in HEK 293Ts (Figure 1G-H). In support of these results, coexpression of a hyperactive catalytic subunit of PI3 kinase mutant (H1056R) with wild type MASTL in HEK293 T cells degraded MASTL protein levels to similar extent as that of wild type AKT (Figure 1I-J). From these findings, it was confirmed that MASTL protein degradation was caused by AKT.

**Figure 1.**
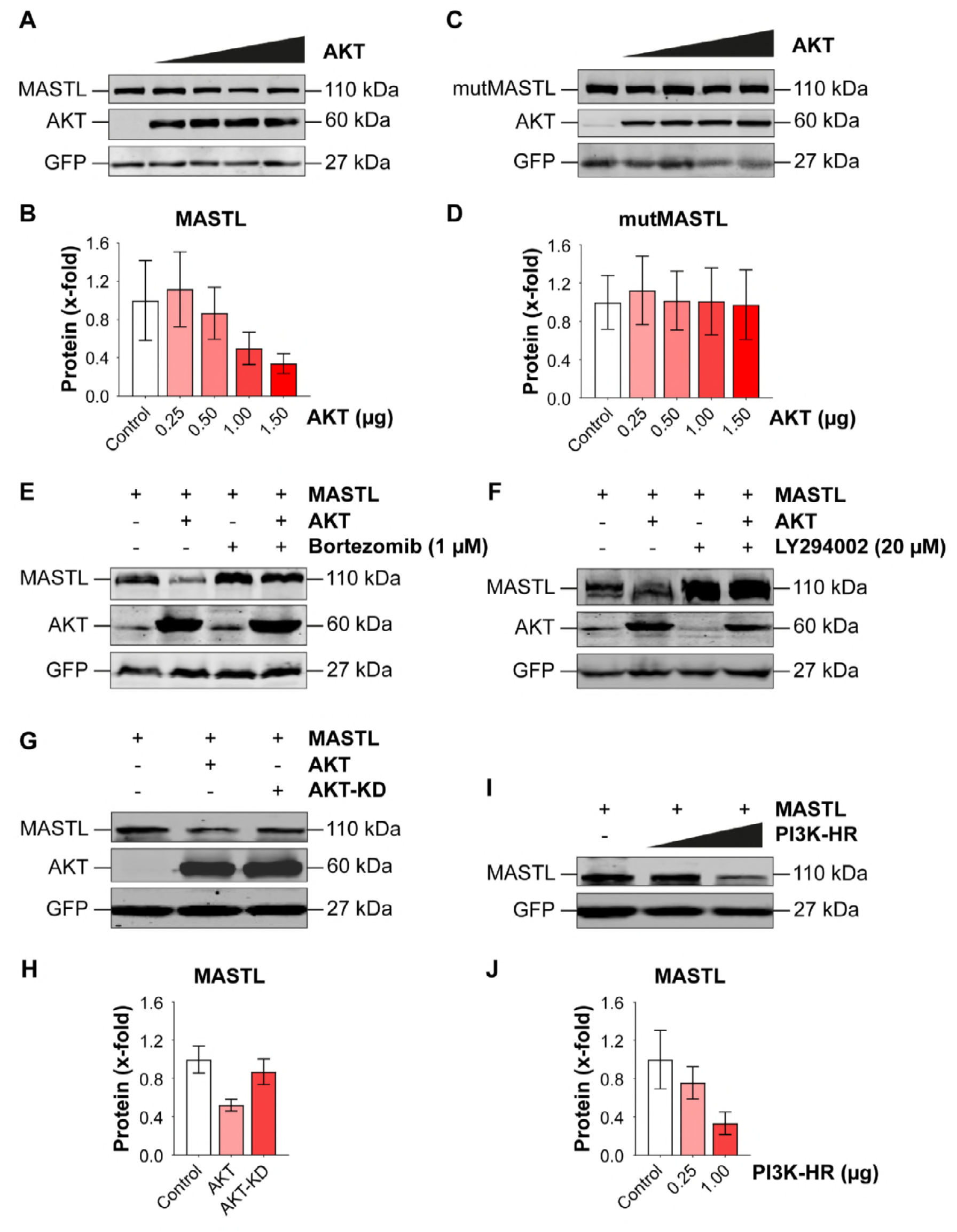
MASTL is a potential target for AKT phosphorylation. MASTL was overexpressed along with increasing expression of AKT using pCMV-HA-MASTL and pCDNA3.1-AKT constructs in HEK 293T cells. GFP construct was used as a transfection control. A) Blot showing MASTL protein levels after adding increasing amounts of AKT DNA. C) MASTL mutant (T299A) did not show any change in expression after addition of AKT DNA. E) Western blot showing MASTL protein levels in presence of AKT after addition of 1μM bortezomib for 24 hours. F) Blot showing proteasomal degradation of MASTL caused by AKT (lane 2 from left) was rescued by inhibiting AKT by the addition of 20μM LY294002 for 18 hours (lane 4). G) Western blot showing the degradation of MASTL protein in presence of wild type AKT was not observed in case of kinase dead mutant of AKT (KD). I) The hyperactive mutant of PI3 Kinase H1056R construct when co-transfected with pCMV-HA-MASTL showed effect similar to that of wild type AKT. B, D, H and J demonstrate the quantification analysis of the respective blots. The experiments were repeated three times. Data is expressed as mean ±SEM of triplicates.

### MASTL is phosphorylated by AKT on its substrate site

The proteasomal degradation of MASTL in presence of AKT implied that MASTL might be phosphorylated by AKT. Therefore, our next aim was to directly confirm whether AKT phosphorylates MASTL on its substrate site. pCMV-HA-MASTL and pCDNA3.1-HA-AKT were co-transfected in HEK 293T cells and untransfected cells were taken as control. This was followed by immunoprecipitation of exogenously expressed HA-MASTL using anti-HA antibody. The immunoprecipitated proteins were blotted with anti phospho-AKT substrate antibody. This antibody recognizes the potential substrates that are phosphorylated by AKT on its consensus site (RXRXXS/T). As can be seen from Figure 2A, the phosphorylation signal for AKT substrate is very strong in MASTL/AKT co-transfected lane as compared to the MASTL lane (Figure 2A). The same blot was reprobed using anti-HA antibody in order to confirm the presence of MASTL and AKT in all the lanes. (Figure 2B). To further confirm whether the phosphorylation was done by AKT specifically at its substrate site on MASTL, AKT was co-transfected with mutant MASTL (T299A) as well as wild type MASTL. After transfection, protein lysates were again subjected to immunoprecipitation using anti HA antibody to pull down HA-MASTL protein. Following Western blotting, the blot was probed using anti-phospho AKT substrate antibody. Our results clearly showed that anti-phospho AKT antibody specifically detected wild type MASTL but not the mutant MASTL (T299A) when co-transfected with AKT (Figure 2C). The same blot was reprobed with anti-HA antibody to detect the presence of other transfected proteins (Figure 2D).

**Figure 2.**
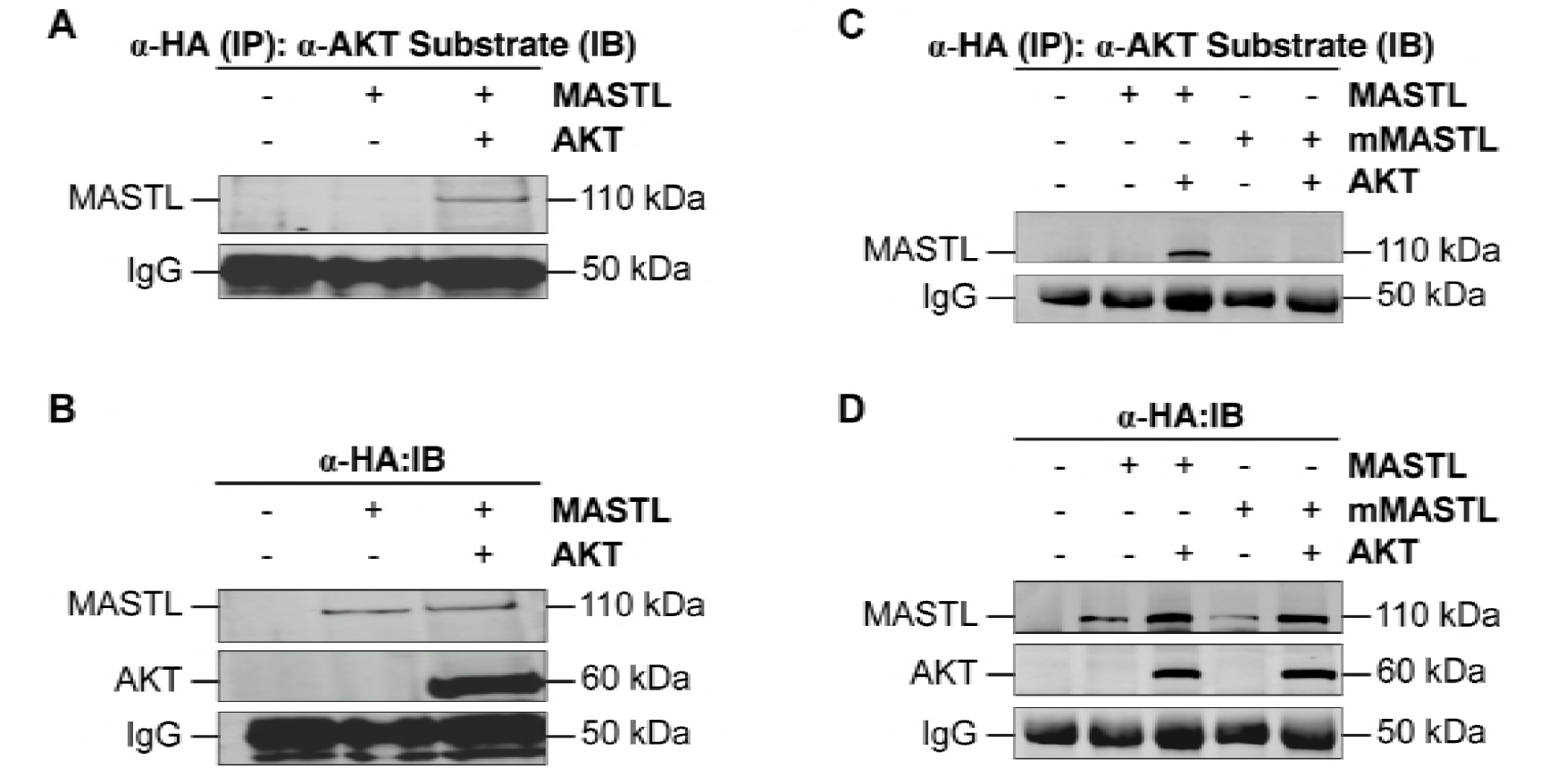
MASTL is phosphorylated by AKT on its substrate site. A-B) pCMV-HA-MASTL was cotransfected with pCDNA3.1-AKT in HEK 293T cells, keeping untransfected cells as control. Cell lysates were subjected to immunoprecipitation by anti HA antibody followed by Western blotting and detected with anti-phospho AKT substrate antibody or HA antibody (after reprobing). C) The same experiment was repeated with mutant MASTL T299A (mMASTL). D) Detection of all the transfected proteins in the same blot after probing it with anti HA antibody.

### AKT phosphorylates MASTL and increases CDK1 substrate phosphorylations during mitotic progression

From the above results, it was confirmed that AKT phosphorylates MASTL at T299 residue. Since MASTL is a mitotic protein and is required in its active form only during mitosis, so we asked whether MASTL would be phosphorylated by AKT only during mitosis. To check that, we co-transfected MASTL and AKT in HEK 293T cells and arrested them in mitosis using 50nM nocodazole for 18 hours. Western blotting results indeed confirmed that AKT phosphorylates MASTL during mitotic phase as was indicated by a distinct band shift on MASTL (Figure 3A).

**Figure 3.**
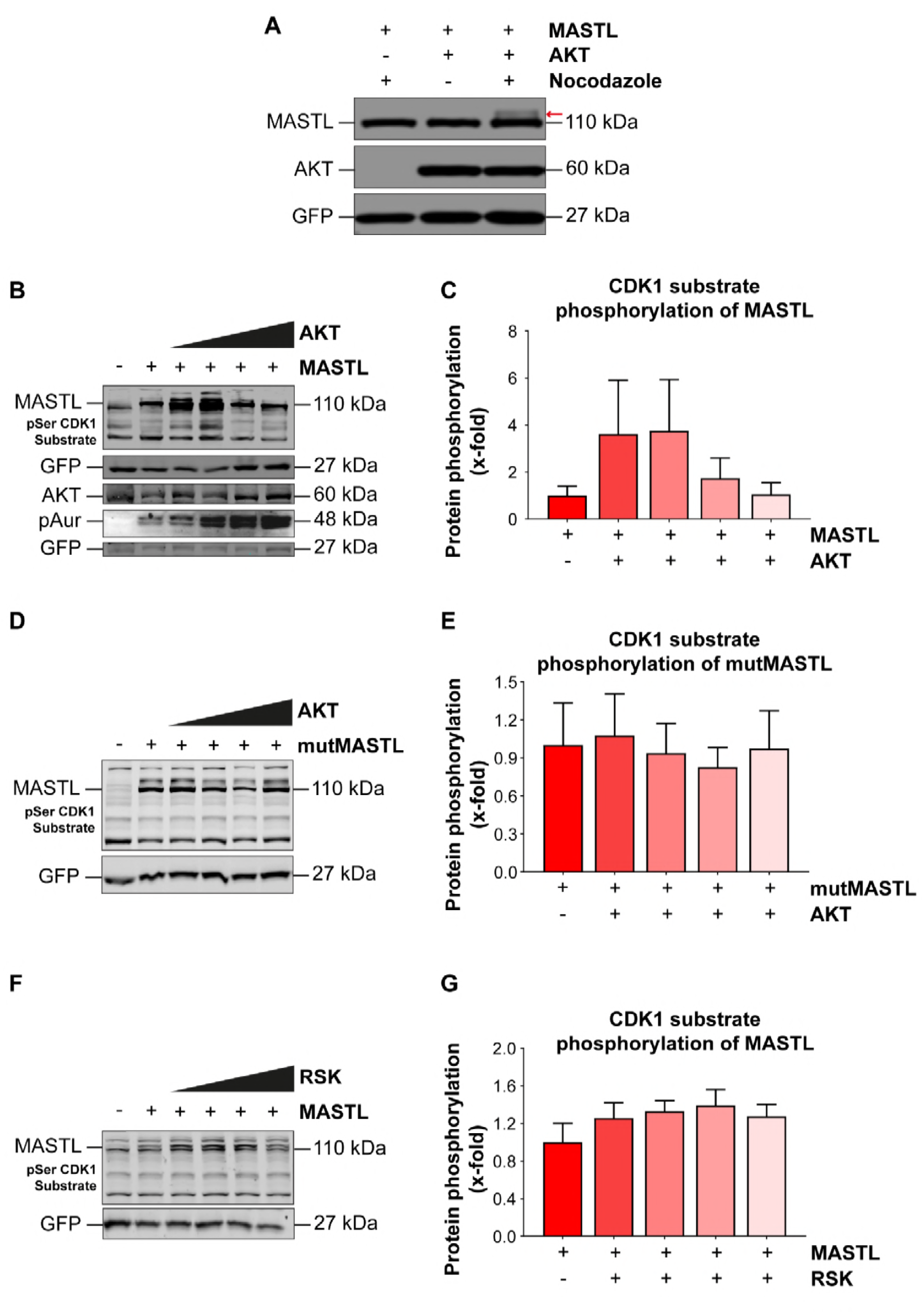
AKT mediated phosphorylation on MASTL increases CDK1 substrate phosphorylations. A) pCMV-HA-MASTL was co-transfected with pCDNA3.1-AKT in HEK 293T cells and GFP was cotransfected as a transfection control. 24 hours post transfection, cells were arrested in mitosis using 50nM nocodazole for 18 hours. Western blot showing the band shift of MASTL in presence of AKT (indicated by red arrow). B) Increasing amounts of pCDNA3.1-AKT were co-transfected with pCMV-HA-MASTL in HEK 293T cells. Blot showing CDK substrate phosphorylation of MASTL as well as other CDK substrates and phosphorylation levels of Aurora kinase with increasing expression of AKT. D) Western blot showing CDK mediated phosphorylations of mutant MASTL or other CDK substrates in presence of increasing amounts of AKT. F) Blot showing CDK1 substrate phosphorylation levels in presence of wild type MASTL and RSK. C, E, G shows the quantification of the respective blots. The experiments were repeated three times. Data is expressed as mean ±SEM of triplicates.

Next, we were interested to determine the effect of AKT phosphorylation on MASTL activation. If AKT mediated phosphorylation at T299 activates MASTL, then cotransfecting MASTL and AKT in cells should increase the percentage of cells in mitosis. Since MASTL activates CDK1 and its substrates by inhibiting PP2A [6], so we used the levels of CDK substrate phosphorylations as mitotic marker in MASTL/AKT cotransfected cells. Increasing amounts of AKT starting from 0.25μg to 1.5μg were cotransfected with MASTL in 293T cells. The extracted lysates were subjected to Western blotting and analyzed for CDK substrate phosphorylation content using anti phospho-CDK substrate antibody. Our results showed that increasing amounts of AKT when cotransfected with MASTL resulted in increased CDK mediated phosphorylations of MASTL as well as other substrates of CDK1 (Figure 3B-C). In addition, phosphorylation levels of Aurora kinase which is used as mitotic markers in cells, also showed an impressive increase in these samples. In contrast to the wild type MASTL, AKT was not able to increase CDK1 phosphorylations in presence of mutant MASTL (Figure 3D-E). These results clearly confirmed that AKT mediated phosphorylation of MASTL is necessary for its activation. Furthermore, a study on the regulation of MASTL have shown that besides CDK1, MASTL is also probably phosphorylated and activated by one of the AGC kinase family members, ribosomal S6 kinase (RSK) [17]. This family of kinases includes AKT as well as RSK, and all members have the same substrate motif for phosphorylation (RXRXXS/T). We wanted to compare the activity of RSK and AKT towards the phosphorylation of MASTL. Our results showed that co-transfection of RSK with MASTL in HEK 293T cells had a very modest effect if at all on CDK substrate levels even at higher amounts of RSK transfected into these cells (Figure 3F-G). Hence these results confirmed that AKT instead of RSK is a bona fide and potent regulator of MASTL.

It was also observed from Figure 3B that at higher concentration of AKT (1*μ*g), CDK phosphorylations of both MASTL as well as other CDK substrates were drastically decreased. This was probably due to high rate of CDK1 mediated phosphorylations of its substrates at increased amount of transfected AKT, followed by proteasomal degradation. To see whether this was true, we co-transfected MASTL and AKT (1*μ*g) in HEK 293T cells and added 1μM bortezomib for 24 hours to inhibit proteasomal degradation. From the results it was clear that bortezomib could prevent the degradation of MASTL and other substrates of CDK in presence of over expressed AKT (Figure 4A). To gain further support for the role of AKT in CDK substrate phosphorylations, we cotransfected kinase dead mutant of AKT (KD) along with MASTL in HEK293T cells. Western blotting results clearly indicated that, unlike the wild type AKT, kinase dead mutant of AKT had no impact on CDK substrate phosphorylations even at higher amounts (Figure 4B). In support of these results, co-transfection of constitutively active mutant of PI3kinase (PI3K 1056HR) with MASTL in HEK 293T cells showed the same effect as that of AKT (Figure 4C).

**Figure 4.**
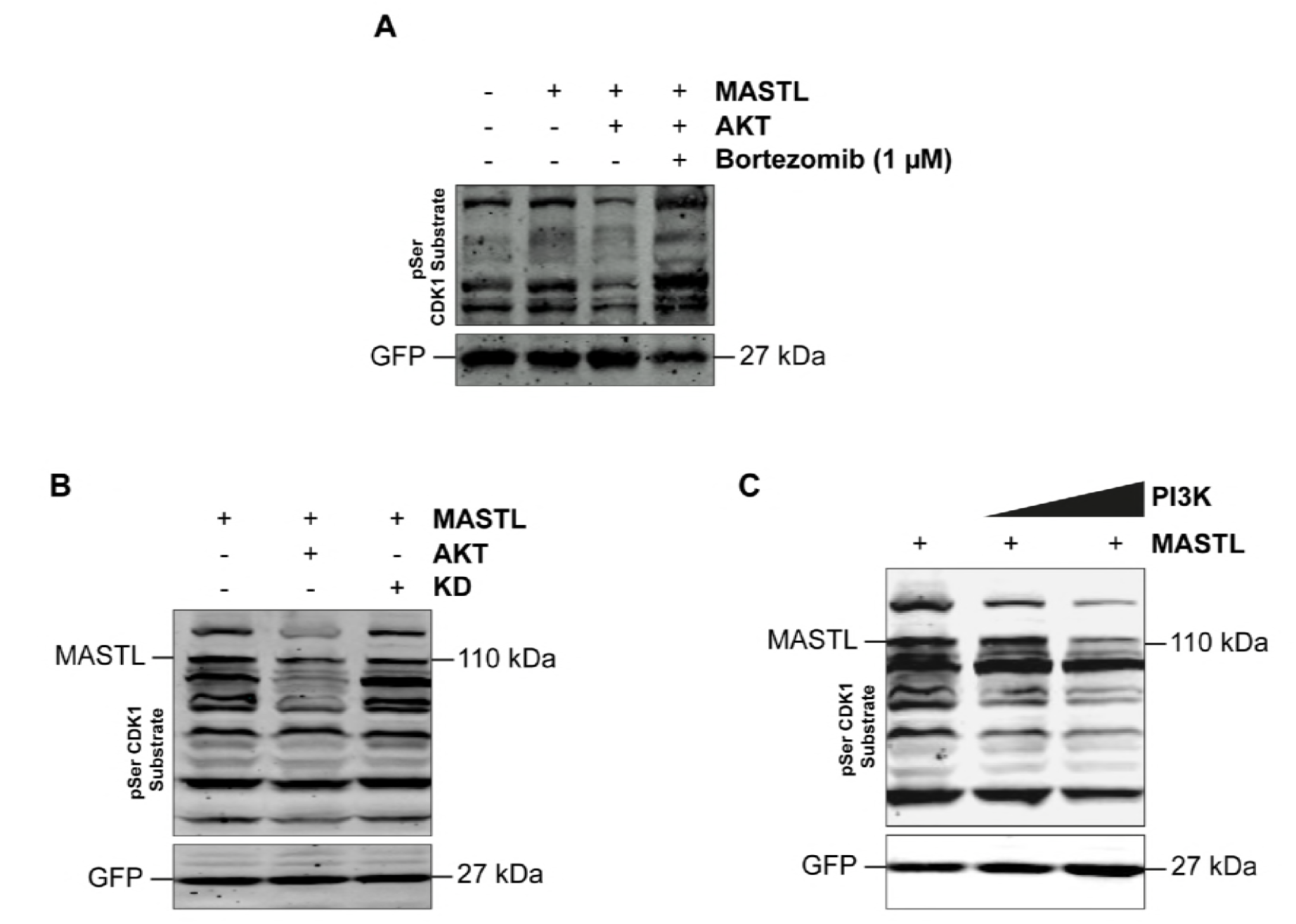
Proteosomal degradation of MASTL and other CDK1 substrates is mediated by AKT. A) MASTL and AKT were over expressed in HEK 293Ts as indicated and 24 hours post transfection, cells were treated with 1μM bortezomib for 24 hours and blotted with anti CDK1 substrate antibody. B) CDK substrate phosphorylations in HEK 293Ts after co-transfection of MASTL with wild type AKT or the kinase dead mutant of AKT (KD). C) Effect of hyperactive mutant of the catalytic subunit p110α of PI3K (1057HR) co-transfected with MASTL on CDK substrate phosphorylation levels in HEK 293Ts.

### AKT promotes cell growth through MASTL

To see whether AKT promotes the cell growth and proliferation of cells through MASTL, we first examined MASTL expression in various cell lines (Figure 5A). Among these cell lines SW480 and HCT116 colorectal cell lines showed highly up-regulated expression of MASTL. We selected SW480 cell line for our further studies since it had comparatively higher expressionof MASTL than in HCT-116 cell line. We stably transfected myristoylated Flag-AKT, shMASTL and two different shAKT constructs (shAKT1 & shAKT2) and checked the expression of these proteins by Western blotting (Figure 5 B-E). To test the effect of AKT on the growth of SW480 cell line, we seeded equal number of these cell lines in 6 well plates, keeping one of the plates as a control. The cells were seeded in triplicates and allowed to grow until they reached approximately 60% of confluence. At this stage, shMASTL and shAKT constructs were induced using 20*μ*g/ml doxycycline for 24 hours. To show that AKT and MASTL promote cell proliferation, we performed crystal violet staining and MTT growth assay (Figure 5, F-H). As can be seen from the figure, AKT overexpressing SW480 cell line showed higher intensity of crystal violet staining (Figure 5F-G) which was also evident from increased cell growth of this cell line observed in MTT assay as compared to control SW480 cell line (Figure 5H). The silencing of MASTL using shMASTL substantially reduced the growth rate of both AKT overexpressing as well as control SW480 cell line. In AKT overexpressing SW480 cell line, the silencing of MASTL resulted in the killing of SW480 cancerous cells comparable to silencing of AKT. In order to see whether the increase in cell proliferation of SW-480 AKT cell line was due to increase in mitosis, we subjected SW-480, SW-480 AKT, SW-480 shMASTL and SW-480 AKT shMASTL cell lines to cell cycle analysis by FACS. We observed an aproximately 10% increase in mitotic population of cells (G2/M peak as seen in FACS analysis) in SW-480 AKT cell line as compared to SW480 control. However, the increase in mitotic index was almost fully reversed by silencing of MASTL in SW480-AKT cell line (Figure 6 A-E).

**Figure 5.**
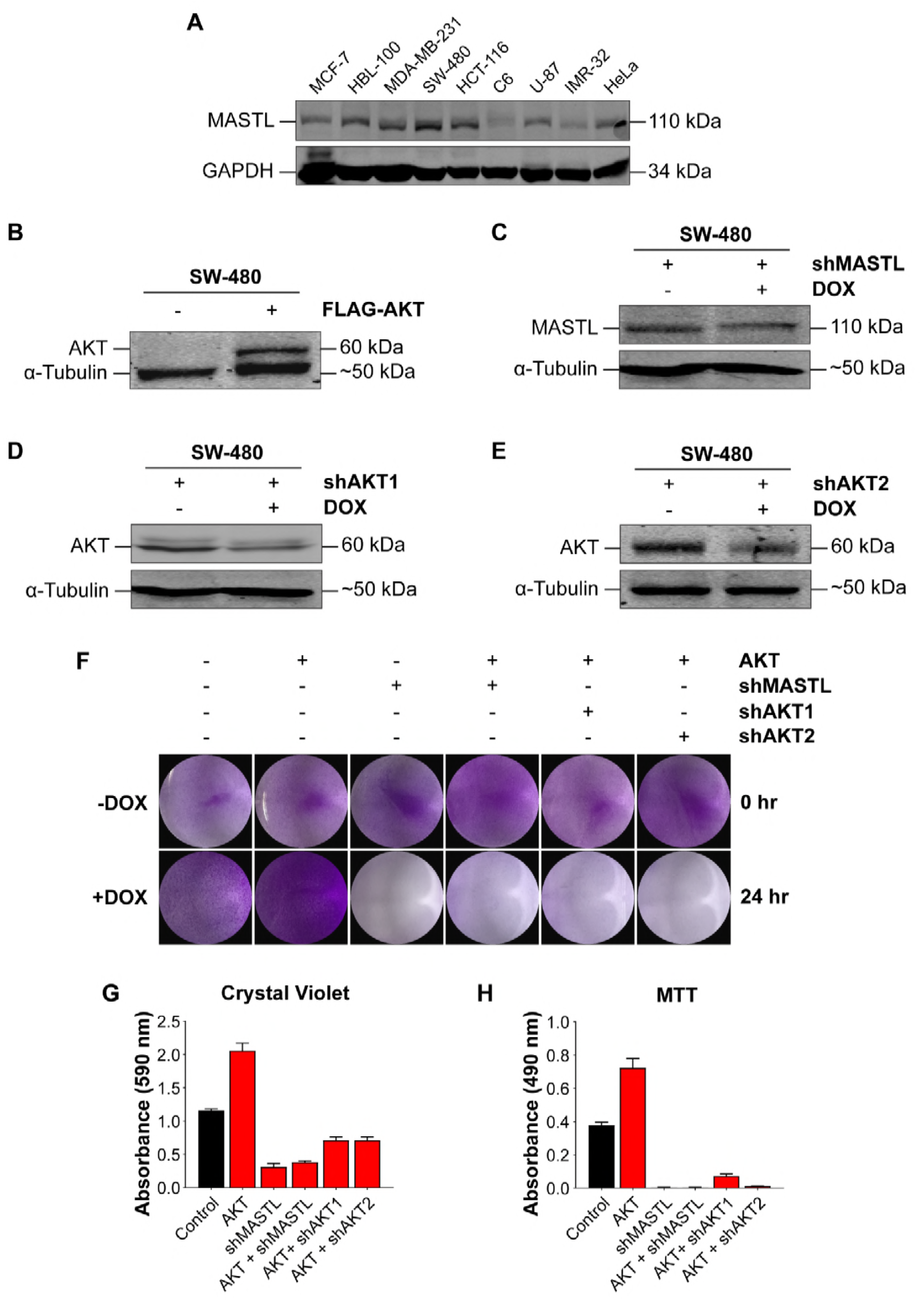
Silencing of MASTL is sufficient to overcome the growth of AKT overexpressing SW480 cell line. A) Various cancerous cell lines showing endogeneous expression of MASTL. SW480 cells had the highest MASTL expression among these cell lines. B) Western blot of SW480 cell extracts showing expression of stably transfected myristilated FLAG-AKT. C-E) Western blot of inducible PLKO-0.1-shMASTL, shAKT 1 & 2 stable SW480 colorectal cell line extracts respectively, showing silencing of each construct after induction by 20μg/ml doxycycline addition. F-H) Cell growth and proliferation assays of SW480 stably expressing AKT, shMASTL, shAKT1 and shAKT2. F-G) Crystal violet staining; H) Cell proliferartion by MTT assay.

**Figure 6.**
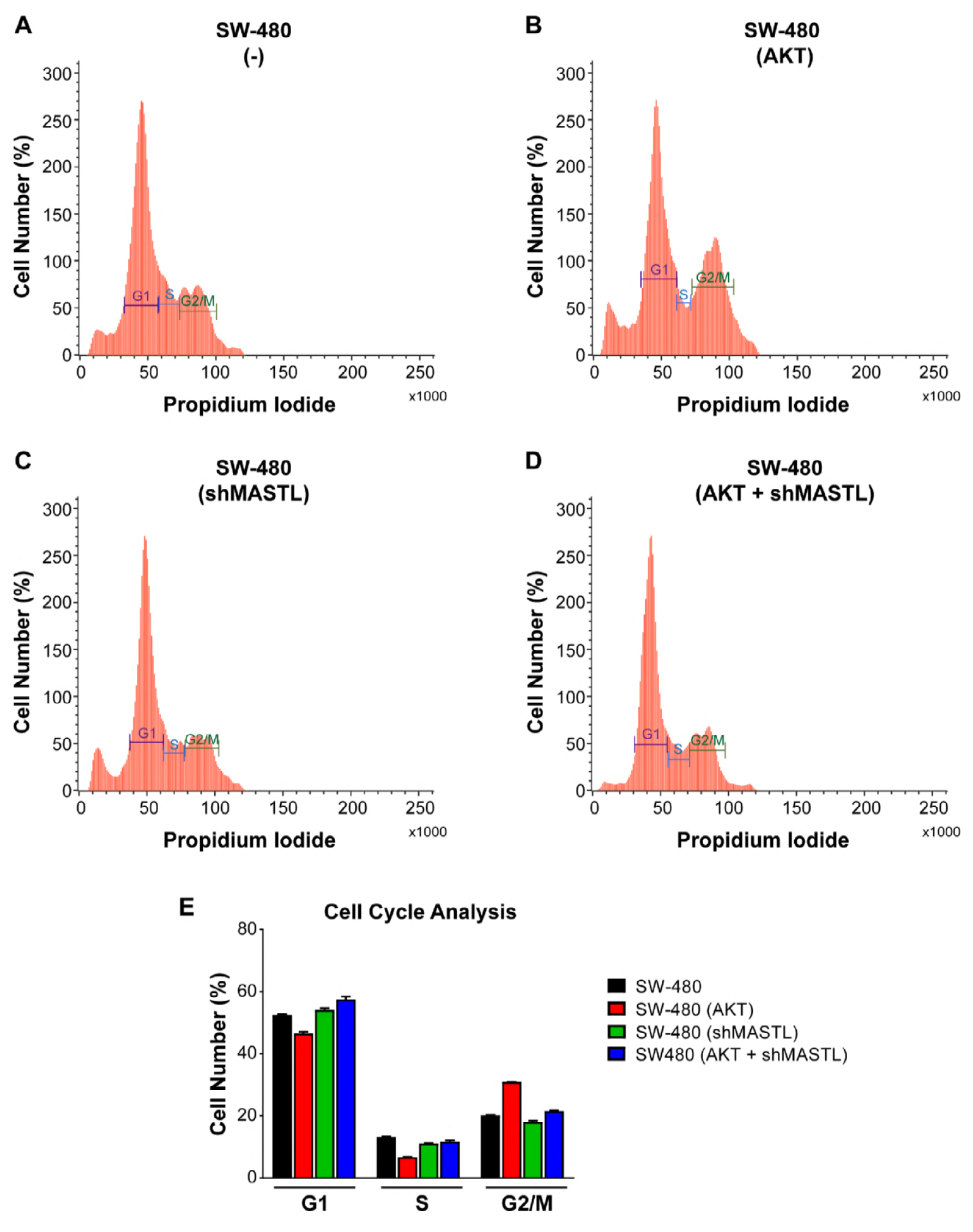
Cell cycle analysis using FACS. A) SW-480 B) SW-480 AKT C) SW-480 shMASTL D) SW-480 AKT shMASTL E) Graphical representation of FACS data obtained from A-D. The increase in G2/M population of cells by AKT overexpression (as in B) was reversed by the knockdown of MASTL in SW-480 cells (as in D).

**Figure 7.**
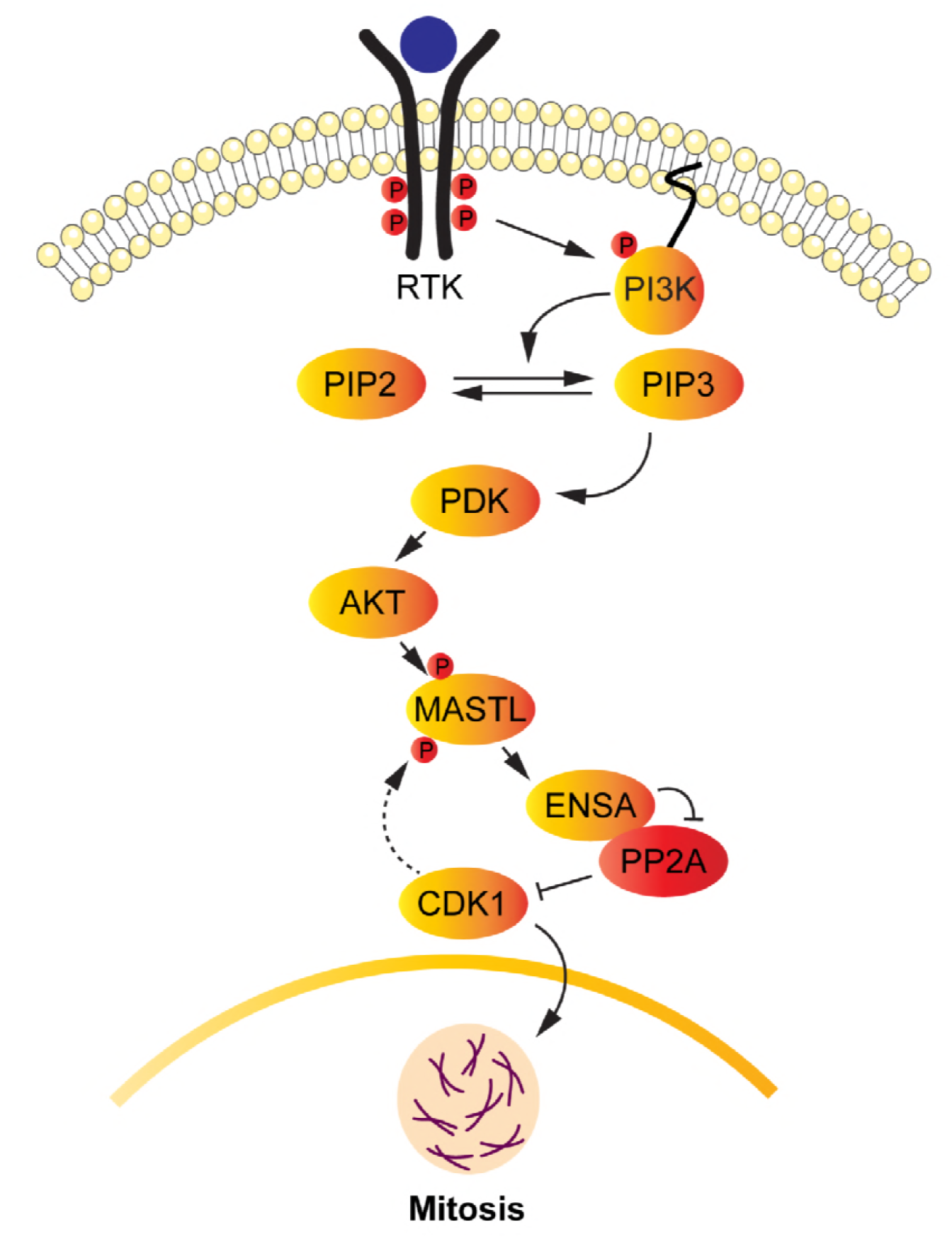
Proposed model showing the connecting link between AKT and MASTL signaling during mitosis. Activation of the PI3 kinase pathway leads to the phosphorylation and activation of AKT. Activated AKT leads to the phosphorylation of MASTL at T299, thereby leading to its activation. Once activated, MASTL leads to the phosphorylation of ENSA which results to the inactivation of PP2A-B55δ. As a result, CDK1/Cyclin B1 complex remains activated, leading to the phosphorylation of numerous substrates including MASTL which plays an important role in the entry and progression of the cells into mitosis.

Altogether, these results suggest that AKT increases cell proliferation of SW480 cell line by enhancing the mitotic progression of cells through the activation of MASTL. And the increase in cell growth of this stably expressing AKT cell line can be significantly reduced by silencing MASTL.

## Discussion

For the continuous proliferation of tumor cells, mitotic proteins are either up-regulated or hyperactivated to promote mitosis [18]. Hence, these proteins have been always found to be interesting and effective targets to overcome cellular proliferation of tumor cells [19]. Microtubule associated serine threonine kinase like (MASTL) is one of the important regulators of mitosis that was discovered a few years ago. The mutated form of MASTL has been seen to be implicated in thrombocytopenia, and up-regulation of this protein has been observed in various cancers [20, 21]. The mechanism of MASTL in cell cycle regulation and cancer progression is still not very well understood. Though there are various studies regarding its role in cell cycle, the understanding of its regulation during mitosis is still poor. There are only a couple of studies that have reported that MASTL is regulated mainly by CDK1 whereas Plx1 can substitute CDK1 by phosphorylating and activating MASTL at the same phosphorylation site at least *in vitro*. The same study has reported that besides CDK1, there might be some other regulators of MASTL potentially from its own AGC family members [22, 23]. We compared the impact of both RSK and AKT on MASTL activity and showed that AKT has more impact on MASTL phosphorylation than RSK, thus suggesting that MASTL could be the bona fide taget for AKT *in vivo*.

In order to better understand the process of cancer progression and developing new therapeutic targets against this disease, studying the regulation and role of new targets like MASTL is of great importance. Therefore, in this study we attempted to elucidate the mechanism of MASTL during mitosis. We employed www.scansite3.edu, an online computational software to search for potential regulators of MASTL and found out that AKT could be one of the regulators of this kinase. Interestingly, MASTL has a perfect consensus site for AKT phosphorylation which is conserved between various mammalian species that prompted us to investigate these findings further. We overexpressed MASTL and AKT together in HEK293T cells and found that MASTL protein is degraded after co-expressing with AKT. Using various chemical inhibitors like bortezomib and LY294002, we observed that the degradation is mediated by proteasomes. Further biochemical assays showed that AKT phosphorylates MASTL on T299 during mitosis. Moreover, AKT mediated phosphorylation of MASTL during mitosis leads to increase in the phosphorylations of CDK1 downstream substrates with a significant increase in phosphorylation of MASTL. It has been observed that various phosphorylations at the N- terminal and C- terminal domain of MASTL lead these two lobes to interact with each other which results in the stabilization of phosphorylations of this kinase [17]. Our results also demonstrate similar mechanism for AKT mediated phosphorylation on MASTL which results in the stabilization of CDK1 mediated phosphorylation of this kinase, leading to its further activation. In contrast, higher expression of AKT leads to the increased degradation of CDK1 substrates. Furthermore, stable expression of AKT in SW480 colorectal cell line that has high expression of MASTL, increased the proliferation of these cells. This increase in SW480-AKT cell growth was reduced by silencing of MASTL as well as AKT in these cells. Moreover, as shown by FACS analysis, AKT enhances the mitotic cell population of SW480 cell line which was seen to be reversed by silencing of MASTL. These findings suggest that overexpression of AKT in SW480 cell line leads to the activation of MASTL which promotes the entry of cells into mitosis.

Our data supports the findings of an earlier report that mentions that GWL leads to the activation of AKT [14]. In this paper, the authors have shown evidence that GWL indirectly leads to the enhanced phosphorylation of AKT at S473, leading to its activation. It does so by an unknown mechanism that stimulates the dephosphorylation and activation of GSK3, which phosphorylates PHLPP at numerous sites, promoting its degradation by the SCF-β-TRCP proteasomal complex. Since PHLPP phosphatase is known to dephosphorylate AKT at S473, hence its degradation by GWL results in the sustained phosphorylation and activation of AKT. Our studies have shown the reciprocal mechanism. We showed evidence that AKT directly phosphorylates MASTL at T299 residue, leading to its activation which leads to the enhanced entry of cells into mitosis. It is possible that both AKT and MASTL reciprocally phosphorylate and activate each other by feed back mechanisms, and thus promote cellular transformation. Though both kinases have been individually seen to be activated in cancers, it will be interesting to see whether both kinases are activated in tumors at the same time. The phosphorylation may occur during a short lived transient interaction between them as we were not successful in finding an interaction between AKT and MASTL in immunoprecipitation experiments (data not shown).

In summary, our findings demonstrate that MASTL is phosphorylated by AKT during mitosis which leads to the further activation of MASTL. Also, elimination of MASTL can overcome uncontrolled cellular proliferation of both MASTL as well as AKT overexpressing cancerous cells. These results assume significance since both AKT and MASTL are reported to be activated and/or overexpressed in many types of cancers and are popular targets for anti-cancer therapy. Since AKT regulates numerous signaling pathways including cell survival pathways, using pan anti-AKT drugs can have toxic effects on the cells and patients. On the other hand, identifying and inhibiting specific downstream candidates like MASTL that regulate mitotic progression may prove to more beneficial as anti-cancer tehrapeutic agents.

## Materials and Methods

### Materials and reagents

Doxycycline, rabbit monoclonal anti-HA, protease and phosphatase inhibitors, polybrene, agarose A/G beads, propidium iodide, RNAase A, MTT, crystal violet, were obtained from Sigma Aldrich. Primers used for PCR were purchased from Integrated DNA technology (IDT). Anti AKT substrate, anti CDK substrate, anti MASTL, anti-Myc (tag), anti GFP, anti phospho Aurora, anti-cyclin B, anti HA and secondary antibodies (DyLight 680 and Dylight 800) for LICOR imaging system were purchased from Cell Signaling Technology, USA. Nitrocellulose membrane, protein ladder, acrylamide solutions and Bradford reagent were purchased from Bio-Rad laboratories USA. Drugs (nocodazole, LY294002, bortezomib) were purchased from Calbiochem/EMD Biosciences. Dulbecco’s Modified Eagle’s Medium (DMEM), fetal bovine serum, penicillin/streptomycin and trypsin were obtained from (Invitrogen/Life Technologies). Q5-high fidelity DNA polymerase, dNTPs, T4 DNA ligase, DpnI and all restriction enzymes that were used in our experiments were purchased from New England Biolabs, USA. Transfection reagent Polyethaneamine (PEI) was obtained from Polysciences, USA. MASTL construct was obtained from Anna Castro, France [22], amplified by PCR using adaptors containing XhoI and NotI restriction sites and was later cloned into pCMV-HA-3.7, using the primer sequences: MASTL (5’-GGCCTCGAGTAGATCCCACCGCGGGAAGCAA-3’ and 3’-GGCCGCGGCCGCCTACAGACTAAATCCAGATAC-5’) by following standard cloning procedures. pCDNA3.1 HA-AKT1 (Addgene plasmid # 9008) [24], PKH3 human RSK (Addgene plasmid # 13841) [25], pCDNA3 T7 AKT1 K179M T308A S473A (Addgene plasmid # 9031) [24]. PI3K (H1056R), HEK293T, Pheonix and SW-480 cell lines were kindly given as gift items by Prof. Thomas Roberts, DFCI, Harvard University, USA. shAKT1 and shAKT2 were cloned in pLKO.1 puro using the following sequences: shAKT1: CGCGTGACCATGAACGAGTTT and shAKT2: CGAGTTTGAGTACCTGAAGCT [26].

### Site directed mutagenesis

For site directed mutagenesis, pCMV-HA-MASTL was used as template to generate the mutant MASTL. Overlapping primers 5’-AGGAAAAGGCTGGCCGCATCCAGTGCCAGTAGT-3’ and 5’-ACTACTGGCACTGGATGCGGCCAGCCTTTTCCT-3’ containing point mutation (A to G at 895^th^ nucleotide of MASTL), were used to amplify the template. This mutation was used to alter the amino acid Serine to Alanine at 299^th^ position on MASTL, corresponding to the AKT phosphorylation site. The protocol from Agilent Quick-change site directed mutagenesis kit was followed to create the mutant. The amplified clones were sent to Scigenome labs, India (www.scigenom.com) for sequencing for the confirmation of the incorporation of the desired mutation.

### Cell culture

Cells were maintained in Dulbecco’s Modified Eagle’s Medium (DMEM) supplemented with 10% FBS and 1% penicillin/streptomycin. Induction of protein expression was done by adding doxycycline at 20μg/mL concentration.

### Transfection

For transient transfections, pCMV-HA-MASTL, pCDNA3.1-HA-AKT, EGFP, PI3K-110*α* (H1056R), pCDNA3T7-AKT1-K179M-T308A-S473A (KD), pKH3RSK, pCMV-HA mutant MASTL were transfected in HEK293T cells using polyethaneamine (PEI) as transfection reagent. The cells were harvested after 48 hours using NP-40 lysis buffer (1% NP40, Tris-Cl 50mM, NaCl 150mM, glycerol 10% and EDTA 2mM) containing protease and phosphatase inhibitors and incubated on ice for about 30 minutes. Denaturation was done by adding 5x loading dye.

The stable cell line of AKT overexpression in SW480 was made by retroviral transfection of 2.5μg of pWZL-Neo-MF-AKT (Addgene USA) along with 2.0μg of vectors expressing gag/pol and vsvg in the ratio of 2:1 in Phoenix cells that were about 80% confluent at the time of transfection. Next day, the medium of the transfected cells was changed and the cells were allowed to grow further for 24 hours. After incubation, the viral supernatant of these transfected phoenix cells along with polybrene 8μg/mL was added to the SW480 cells for at least 6 hours. At the end of the day, cells were washed with 1xPBS and fresh medium was added for overnight and the same procedure was repeated on the next day. On the third day, the cells were washed and the selection of stably transfected cells was started by adding 500μg/mL neomycin to the medium. After selecting the cells for about two weeks, each cell line was amplified and used for further experiments.

### Western Blotting

The denatured cell lysates were run in SDS-PAGE and the proteins were transferred onto nitrocellulose membrane. Following this, the blots were blocked with 5% non-fat dry milk for about 1 hour. After that, corresponding primary antibodies were added to each blot and incubated overnight at 4°C. Subsequently, washing of blots was done three times by 1x TBS-T (NaCl 13.7mM, Tris 20mM; pH 7.4, Tween-20, 0.05%) for 10 minutes each. This was followed by incubation of blots in secondary antibody for about 1 hour. Finally, the blots were again washed three times with 1xTBS-T and subjected to infrared detection system using LICOR machine.

### Immunoprecipitation

Anti-HA antibody was premixed with protein G agarose beads in the ratio of 1:20. The antibody and bead mixture were kept on a rotator for about 1hr at 4°C. The preloaded beads were then washed three times with 1x PBS at 3,000 rpm for 1 minute each. Equal amount of each protein extracts (as measured by Bradford assay) were added to the protein G Agarose beads pre-loaded with primary antibody. The samples were incubated/mixed overnight on a rotator at 4°C. After incubation, the beads were washed three times with 1x PBS at 3,000 rpm for 1 minute each. Finally, 2x loading buffer was added to the beads and the mixtures were boiled at 100°C for 3-5 minutes. The samples were then subjected to Western blotting.

### Crystal violet

Equal number of cells (1×10^5^) were seeded in 6 well plates. Next day, the induction of shMASTL, shAKT1 and shAKT2 with 20μg/mL doxycycline was done for 24 hours after which the plates were taken and washed once with 1xPBS. Following that, cells were fixed with methanol for 10 minutes and washed again with 1xPBS. 1 mL of 0.1% crystal violet (0.1% crystal violet in 25% ethanol) was added per well and left in the wells for about 10 minutes at room temperature. The dye was then aspirated from the plates, followed by washing with distilled water. The stained cells were allowed to dry at room temperature. For quantification, the dye attached to the cells was extracted with 2mL of 10% acetic acid, and its absorbance was measured at 590nm. Experiments were done in duplicates and were repeated three times.

### MTT assay

Equal number of cells (10,000) were plated in 12 well plates. After reaching to 50% cell density, the induction of shMASTL, shAKT1 and shAKT2 with 20*μ*g/mL doxycycline was done for 24 hours. Next, the medium of cells was replaced with fresh medium and 100*μ*L of 12mM MTT stock (5mg MTT in 1mL 1x PBS) was added to each well. After incubating the cells with MTT for about 2 hours in CO_2_ incubator at 37°C, 1mL of SDS-HCI (5 gm of SDS in 50mL 0.01M HCl) solution was added to each well. The solutions were mixed thoroughly by pipetting and incubated again at 37°C for about 4 hours. Next, the plates were removed from the incubator, the solutions were again mixed by pipetting and the absorbance was measured at 570nm.

### FACS analysis

For cell cycle analysis, cell lines were grown up to 60-70% density. At the time of harvesting, cells were washed with 1x PBS (without Ca^2+^ and Mg^2+^). For disaggregation of cells, 3mL of 1xPBS supplemented with 0.1% EDTA was added to the plates and cells were allowed to incubate for 5 minutes in CO_2_ incubator. Cells were then dislodged from the plates by pipetting to obtain single cell suspension. Cells were collected in 15 mL tubes and centrifuged at 1000xg for 5 minutes. Cell pellets were washed with 1xPBS containing 1% serum at 1000xg for 5 minutes. Finally, the cell pellets were resuspended in 0.5mL 1xPBS and fixed by the drop wise addition of about 5 mL chilled ethanol while vortexing to prevent cell clumping. At this stage, the cells were stored at 4°C for at least overnight in the dark. Next day, centrifugation at 1000x g for 5 minutes was done, the pellets were again washed once with 1xPBS/1% serum and resupended in 500*μ*l of propidium iodide/RNase solution (50*μ*g/mL propidium iodide, 10mM Tris, pH7.5, 5mM MgCl2 and 20g/mL RNase A). The resuspended mixture was incubated at 37°C for 30 minutes to remove RNA molecules in the solution. The samples were then subjected to flow cytometric analysis using FACS Verse and analyzed by ModFit software.

## Acknowledgements

We gratefully acknowledge the generous support of the Department of Biotechnology (DBT), Government of India who supported this research project through their grants BT/PR5743/BRB/10/1100/2012, BT/PR13605/MED/30/1525/2015, BT/PR2283/AGR/36/692/2011 and Ramalingaswami Fellowship to Dr. Shaida Andrabi. The support of the Department of Science and Technology (DST), Government of India through their FIST grant to the Department of Biochemistry, University of Kashmir is also gratefully acknowledged. We also thank UGC-CSIR, Government of India for providing fellowships to Irfana Reshi and Misbah Un Nisa, and also DST for supporting S. Qaaifah Gillani, Sameer Bhat and Zarka Sarwar through INSPIRE fellowships.

---

**Figure S1:**
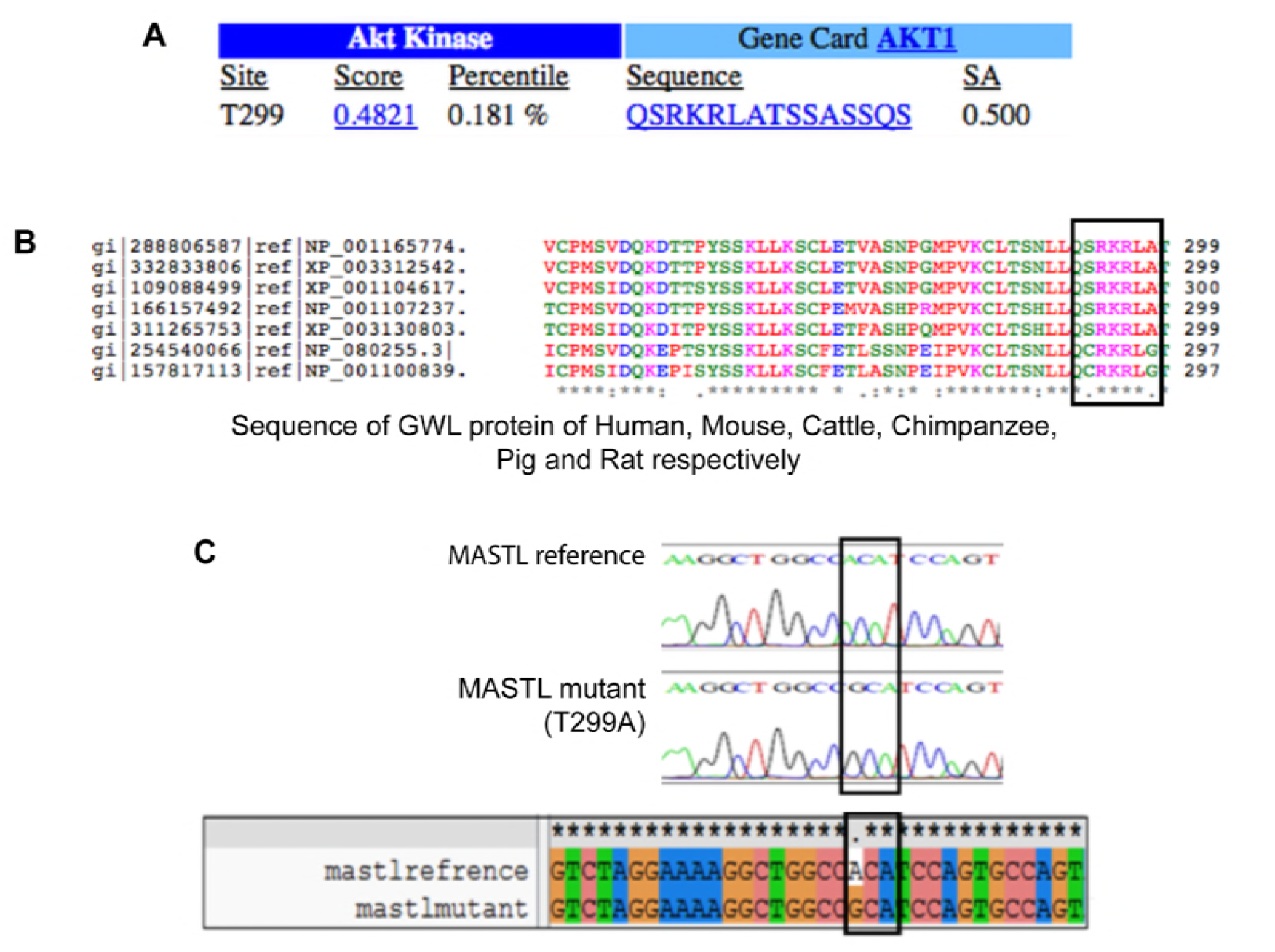
Bioinformatics analysis of MASTL. A) Scan site showing presence of AKT conserverd phosphorylation site (RKRLAT) on MASTL sequence. B) Sequence alignment of MASTL protein from various mammalian species using Clustalw. C) Chromatogram showing confirmation of induction of mutation in MASTL sequence at T299 aminoacid residue.

## References

1. Voets E, Wolthuis RMF: MASTL is the human ortholog of Greatwall kinase that facilitates mitotic entry, anaphase and cytokinesis. Cell cycle (Georgetown, Tex) 2010, 9(17):3591–3601.

2. Yu J, Zhao Y, Li Z, Galas S, Goldberg ML: Greatwall kinase participates in the Cdc2 autoregulatory loop in Xenopus egg extracts. Molecular cell 2006, 22(1):83–91.

3. Mochida S, Hunt T: Protein phosphatases and their regulation in the control of mitosis. EMBO Rep 2012, 13(3):197–203.

4. Mochida S, Ikeo S, Gannon J, Hunt T: Regulated activity of PP2A-B55 delta is crucial for controlling entry into and exit from mitosis in Xenopus egg extracts. The EMBO journal 2009, 28(18):2777–2785.

5. Gharbi-Ayachi A, Labbe JC, Burgess A, Vigneron S, Strub JM, Brioudes E, Van-Dorsselaer A, Castro A, Lorca T: The substrate of Greatwall kinase, Arpp19, controls mitosis by inhibiting protein phosphatase 2A. Science 2010, 330(6011):1673–1677.

6. Mochida S, Maslen SL, Skehel M, Hunt T: Greatwall phosphorylates an inhibitor of protein phosphatase 2A that is essential for mitosis. Science 2010, 330(6011):1670–1673.

7. Barr FA, Elliott PR, Gruneberg U: Protein phosphatases and the regulation of mitosis. J Cell Sci 2011, 124(Pt 14):2323–2334.

8. Voets E, Wolthuis R: MASTL promotes cyclin B1 destruction by enforcing Cdc20-independent binding of cyclin B1 to the APC/C. Biology open 2015, 4(4):484–495.

9. Manchado E, Guillamot M, de Cárcer G, Eguren M, Trickey M, García-Higuera I, Moreno S, Yamano H, Cañamero M, Malumbres M: Targeting mitotic exit leads to tumor regression in vivo: Modulation by Cdk1, Mastl, and the PP2A/B55α, δ phosphatase. Cancer cell 2010, 18(6):641–654.

10. Rogers S, Fey D, McCloy RA, Parker BL, Mitchell NJ, Payne RJ, Daly RJ, James DE, Caldon CE, Watkins DN et al: PP1 initiates the dephosphorylation of MASTL, triggering mitotic exit and bistability in human cells. J Cell Sci 2016, 129(7):1340–1354.

11. Anania M, Gasparri F, Cetti E, Fraietta I, Todoerti K, Miranda C, Mazzoni M, Re C, Colombo R, Ukmar G et al: Identification of thyroid tumor cell vulnerabilities through a siRNA-based functional screening. Oncotarget 2015, 6(33):34629–34648.

12. Alvarez-Fernandez M, Sanz-Flores M, Sanz-Castillo B, Salazar-Roa M, Partida D, Zapatero-Solana E, Ali HR, Manchado E, Lowe S, VanArsdale T et al: Therapeutic relevance of the PP2A-B55 inhibitory kinase MASTL/Greatwall in breast cancer. Cell death and differentiation 2017.

13. Zhuge BZ, Du BR, Meng XL, Zhang YQ: MASTL is a potential poor prognostic indicator in ER+ breast cancer. Eur Rev Med Pharmacol Sci 2017, 21(10):2413–2420.

14. Vera J, Lartigue L, Vigneron S, Gadea G, Gire V, Del Rio M, Soubeyran I, Chibon F, Lorca T, Castro A: Greatwall promotes cell transformation by hyperactivating AKT in human malignancies. eLife 2015, 4.

15. Quaynor SD, Bosley ME, Duckworth CG, Porter KR, Kim SH, Kim HG, Chorich LP, Sullivan ME, Choi JH, Cameron RS et al: Targeted next generation sequencing approach identifies eighteen new candidate genes in normosmic hypogonadotropic hypogonadism and Kallmann syndrome. Molecular and cellular endocrinology 2016, 437:86–96.

16. Wong PY, Ma HT, Lee HJ, Poon RY: MASTL(Greatwall) regulates DNA damage responses by coordinating mitotic entry after checkpoint recovery and APC/C activation. Scientific reports 2016, 6:22230.

17. Vigneron S, Gharbi-Ayachi A, Raymond AA, Burgess A, Labbe JC, Labesse G, Monsarrat B, Lorca T, Castro A: Characterization of the mechanisms controlling Greatwall activity. Mol Cell Biol 2011, 31(11):2262–2275.

18. Perez de Castro I, de Carcer G, Malumbres M: A census of mitotic cancer genes: new insights into tumor cell biology and cancer therapy. Carcinogenesis 2007, 28(5):899–912.

19. Dominguez-Brauer C, Thu KL, Mason JM, Blaser H, Bray MR, Mak TW: Targeting Mitosis in Cancer: Emerging Strategies. Molecular cell 2015, 60(4):524–536.

20. Johnson HJ, Gandhi MJ, Shafizadeh E, Langer NB, Pierce EL, Paw BH, Gilligan DM, Drachman JG: In vivo inactivation of MASTL kinase results in thrombocytopenia. Experimental hematology 2009, 37(8):901–908.

21. Wang L, Luong VQ, Giannini PJ, Peng A: Mastl kinase, a promising therapeutic target, promotes cancer recurrence. Oncotarget 2014, 5(22):11479–11489.

22. Burgess A, Vigneron S, Brioudes E, Labbe JC, Lorca T, Castro A: Loss of human Greatwall results in G2 arrest and multiple mitotic defects due to deregulation of the cyclin B-Cdc2/PP2A balance. Proc Natl Acad Sci U S A 2010, 107(28):12564–12569.

23. Blake-Hodek KA, Williams BC, Zhao Y, Castilho PV, Chen W, Mao Y, Yamamoto TM, Goldberg ML: Determinants for activation of the atypical AGC kinase Greatwall during M phase entry. Mol Cell Biol 2012, 32(8):1337–1353.

24. Ramaswamy S, Nakamura N, Vazquez F, Batt DB, Perera S, Roberts TM, Sellers WR: Regulation of G1 progression by the PTEN tumor suppressor protein is linked to inhibition of the phosphatidylinositol 3-kinase/Akt pathway. Proc Natl Acad Sci U S A 1999, 96(5):2110–2115.

25. Richards SA, Dreisbach VC, Murphy LO, Blenis J: Characterization of regulatory events associated with membrane targeting of p90 ribosomal S6 kinase 1. Mol Cell Biol 2001, 21(21):7470–7480.

26. Vasudevan KM, Barbie DA, Davies MA, Rabinovsky R, McNear CJ, Kim JJ, Hennessy BT, Tseng H, Pochanard P, Kim SY et al: AKT-independent signaling downstream of oncogenic PIK3CA mutations in human cancer. Cancer Cell 2009, 16(1):21–32.

